# Defining and rescuing pathomechanisms of myotubularin and autophagy disruption in a novel human cell model of Charcot-Marie-Tooth Type 4B3

**DOI:** 10.64898/2026.02.03.703518

**Authors:** Malika Sharma, Xinyi S. Mao, Sarah C. Stumpf, Lan Wang, Jason P. Chua

## Abstract

Charcot-Marie-Tooth Type 4B3 (CMT4B3) is a genetic disorder leading to peripheral axon degeneration and clinical manifestations of distal weakness and gait impairment. CMT4B3 is caused by mutations in *SBF1*/MTMR5, a negative regulator of phosphoinositide signaling and autophagy. Although *SBF1* mutations are ubiquitously expressed, how and why loss of *SBF1*/MTMR5 exerts deleterious effects predominantly in neurons of the peripheral nervous system (PNS) is unknown. To investigate the effects of mutant *SBF1*/MTMR5 on PNS neurons compared to non-neurons, we engineered a novel and unique model system of CMT4B3 using human induced pluripotent stem cells (iPSCs) differentiated into key components of the PNS: motor neurons (iMNs), sensory neurons (iSNs), or skeletal muscle (iMuscle). To model CMT4B3, we used iPSCs derived from a CMT43B patient, or genetically knocked down *SBF1* in WT cells. Strikingly, CMT4B3 iMNs showed the highest degree of cell degeneration among all cell types, concordant with the clinical phenotype of patients. We also found that CMT4B3 iMNs and iSNs showed attenuated expression of MTMR5 and related paralogs MTMR2 and MTMR13. Knockdown of *SBF1* most significantly augmented autophagy in iMNs than other cell types. Finally, we tested treatment with VPS34-IN1, a pharmacologic inhibitor of the Class III PI3-Kinase functioning in opposition to MTMR5 in regulating phosphoinositides, and found that VPS34-IN1 rescued cell death in CMT4B3 iMNs. Together, our results for the first time confirm PNS cell type-specific differences in myotubularin expression, autophagy, and vulnerabilities to *SBF1* mutations, and identify a novel therapeutic strategy of high disease-modifying potential for CMT4B3.

## INTRODUCTION

Charcot-Marie-Tooth disease type 4B3 (CMT4B3) is a rare, hereditary, and incurable neurological disorder with heterogenous manifestations^1^. Symptoms include progressive weakness, muscle atrophy, foot deformities, contractures, and gait impairment starting in childhood and adolescence^1-10^. Unique to CMT4B3 and distinct from other subtypes of Charcot-Marie-Tooth, cases of CMT4B3 from Western Asia demonstrate fork and bracket syndrome (referencing characteristic MRI features of cranial nerve degeneration in the pons and midbrain) ^3-7^. Clinically, these patients manifest with microcephaly, cognitive impairment, syndactyly, and multiple cranial neuropathies leading to facial weakness, oculomotor palsies, hearing loss, and dysphagia^3-7^. The most common form of CMT4B3 is an axonal neuropathy, in which electrodiagnostic studies reveal decrements in compound muscle action potentials and sensory amplitudes, and nerve biopsies confirm sensorimotor peripheral nerve loss^3,4,6,8-11^. In the single dysmyelinating variant reported, electrodiagnostic studies show prolonged sensorimotor latencies that correlate with Schwann cell abnormalities of myelin thinning, excessive and redundant myelin folding, and onion bulbs^2^. There is currently no established disease-modifying therapy for CMT4B3, and treatment is entirely supportive^1^.

CMT4B3 is caused by autosomal recessive mutations in *SBF1* (<u>S</u>ET-<u>B</u>inding <u>F</u>actor 1), which encodes the protein MTMR5 (myotubularin-related phosphatase 5)^2^. MTMR5 is a catalytically inactive pseudophosphatase belonging to the 14-member family of myotubularin-related phosphatases^12,13^. Despite lacking a catalytically active phosphatase domain, MTMR5 interacts with and positively regulates its active phosphatase paralog, MTMR2^14,15^. When homodimerized or heterodimerized with MTMR5 or MTMR13, MTMR2 dephosphorylates phosphoinositides (PtdIns), which are membrane lipids that serve as second messengers in activating the autophagy pathway, among other functions^12,14-17^. In line with the molecular connections among MTMR5, MTMR2, and MTMR13, mutations in the genes *MTMR2* and *SBF2* (MTMR13) underlie clinical connections with CMT4B3 by causing the hereditary neuropathies CMT4B1^18^ and CMT4B2^19^, respectively.

Pathogenic mutations in *SBF1* occur throughout the gene and lead to missense amino acid substitutions, deleterious truncations, or transcript splicing defects, and ultimately loss of MTMR5 function^2,6,8-11,20^. Given the role of MTMR5 in metabolizing PtdIns lipids and regulating autophagy through MTMR2, these myotubularins likely act as autophagy suppressors^12,17,21^. In line with this, we previously identified a neuron-selective enrichment of MTMR5 in human induced pluripotent stem cell (iPSC)-derived neurons (iNeurons)^22^. This expression pattern inversely correlated with sensitivity to Torin1-induced autophagy induction, with MTMR5-rich iNeurons the least sensitive, while MTMR5 overexpression suppressed autophagy induction, and MTMR5 knockdown sensitized iNeurons to Torin1-mediated induction of autophagy^22^. Given these results and the inhibitory function attributed to MTMR5, pathogenic mutations in *SBF1*/MTMR5 may lead to toxic over-activation of autophagy in the PNS. Partially in line with this, mitophagy, but not non-selective macroautophagy, appears to be upregulated in CMT4B3 patient fibroblasts^23^. However, these findings have yet to be confirmed in CMT4B3 lower motor and peripheral sensory neurons, which are the cells primarily affected by the disease in human patients^1-5,8,10^.

As a result, the MTMR5-related effects on autophagy and viability in specific cell types of the nervous system, and the reasons why *SBF1*/MTMR5 mutations exert disproportionate toxicity in peripheral neurons, have remained poorly defined. Insights and relevance from prior study results are mixed due to the use of non-neuronal or immortalized cell lines^14,23,24^. Murine knockout of *Sbf1* has provided insights into roles of Mtmr5 in radial axon sorting and spermatogenesis, but mouse modeling serves as a poor surrogate for studying CMT4B3 due to a failure to recapitulate the axonopathy of human patients^25,26^. Without more precise disease modeling, the applicability of preclinical research will be severely limited. Investigating molecular details of CMT4B3 in relevant human cells and cell types is thus paramount for informing effective precision medicine by allowing assessments for maximal rescuing effects in the cell types of greatest vulnerability and that have a relevant, human biological background.

To address these gaps, we engineered a novel human peripheral nerve cell model system of CMT4B3 derived from WT and patient-derived iPSCs. In these cells, we completed stable genomic knock-in of inducible transcription factor cassettes that enabled differentiation into PNS cell types in a rapid and robust manner (<14 days, >99% efficiency). In this way, we were able to generate essentially pure populations of human motor neurons (iMNs), sensory neurons (iSNs), and skeletal muscle cells (iMuscle). All cell types with a given genotype have isogeneity with each other, having derived from a common parental iPSC line, and thus allow direct head-to-head comparisons regarding CMT4B3 pathobiology in unprecedented fashion. With PNS cells harboring WT and CMT4B3 genetic backgrounds in hand, we unraveled cell type-specific and CMT4B3-dependent differences and dysfunction in MTMR expression, autophagy, and cell viability. We thus confirmed the powerful capability of our system to delineate new insights about myotubularin biology and CMT4B3 pathogenesis in the peripheral nervous system.

## MATERIALS & METHODS

### iPSC culture and maintenance

Human induced pluripotent stem cells (iPSCs) were maintained under feeder-free conditions in TeSR-E8 (STEMCELL Technologies) on vitronectin-coated plates, with daily media changes. The CMT4B3 patient-derived iPSC line was generated from skin fibroblasts of an individual carrying compound heterozygous *SBF1/*MTMR5 mutations (maternal: c.3493_3494dupTA; paternal: c.5474_5475delTG). The KOLF2.1J iPSC line^31^ served as the wild-type (WT) control. An additional WT iPSC line expressing mEGFP-LC3B (CRISPR–Cas9 knock-in 5’ to the N-terminal exon of the *MAP1LC3B* gene; Allen Institute for Cell Science, www.allencell.org) was used for live-cell imaging experiments.

### Genomic modification of iPSCs

Tet-ON transcription factor (TF) cassettes were stably integrated into the genomes of iPSCs using a previously established PiggyBac transposase system^28^. Separate cassettes encoded lineage-specific TFs: *ISL1, LHX3*, and *NGN2* for motor neurons (iMNs); *NGN2* and *BRN3a* for sensory neurons (iSNs); and *MYOD1* with *OCT4* shRNA for skeletal muscle cells (iMuscle), and cassettes co-express blue fluorescent protein (BFP) as a selection marker. Medium-sized iPSC colonies were dissociated using EDTA and seeded on vitronectin-coated plates. The next day, transfection was performed with Lipofectamine-STEM (Thermo Fisher Scientific) and equal amounts of PiggyBac and transposase plasmid DNA. After overnight incubation, media were replaced with fresh TeSR-E8, and BFP expression was used to identify and isolate positive clones expressing each Tet-ON cassette, with manual selection and passaging to generate iPSC colonies with 100% clonal populations of cells expressing BFP.

### Differentiation of IPSCs into PNS cells

After clonal genomic integration of Tet-ON TF cassettes, iPSCs were differentiated into iMNs, iSNs, and iMuscle cells using established protocols of doxycycline supplementation^22,30,50^. For each lineage, iPSCs were dissociated with Accutase and plated on PEI-coated or Matrigel-coated plates and slides (iMNs, iSNs), or vitronectin-coated plates and slides (iMuscle), in TeSR-E8 supplemented with 10µM Y-27632 and 2µg/mL doxycycline. For iMNs, cells were cultured with serial media changes as follows: DIV1-3 with TeSR-E8 supplemented with 1X N-2 supplement (Thermo Fisher), 2µg/mL doxycycline (Sigma), and 0.2µg/mL Compound E (Sigma); DIV3-6 with DMEM/F12 (Thermo Fisher) supplemented with 1X N-2 supplement, 1X non-essential amino acids (NEAA), 1X GlutaMAX (Thermo Fisher), 2µg/mL doxycycline, and 0.2µg/mL Compound E; DIV6-14 with Neurobasal-A (Thermo Fisher) supplemented with 1X B-27 (Thermo Fisher), 1X GlutaMAX, 2µg/mL doxycycline, 10ng/mL human BDNF (Peprotech), 5ng/mL NT-3 (Peprotech), and 0.2µg/mL laminin (Sigma), with media top off on DIV10. For iSNs, cells were cultured with serial media changes as follows: DIV1 with TeSR-E8 supplemented with 1X N2, 1X NEAA, 2µg/mL doxycycline, 10ng/mL human BDNF, 5ng/mL NT-3, and 0.2µg/mL laminin; DIV2 with 1:1 mix of TeSR-E8 and DMEM/F12 supplemented with 1X N2, 1X NEAA, 2µg/mL doxycycline, 10ng/mL human BDNF, 5ng/mL NT-3, and 0.2µg/mL laminin; DIV3-14 with Neurobasal-A supplemented with 1X B-27, 1X GlutaMAX, 2µg/mL doxycycline, 10ng/mL human BDNF (Peprotech), 5ng/mL NT-3, and 0.2µg/mL laminin, with media top offs on DIV6 and DIV10. For iMuscle, cells were cultured with serial media changes as follows: DIV1-5 in MEM-α (Thermo Fisher) supplemented with 5% KnockOut Serum Replacement (Thermo Fisher), 1X NEAA, 1X GlutaMAX, 1mM sodium pyruvate (Thermo Fisher), 0.1mM 2-mercaptoethanol (Sigma), and 2µg/mL doxycycline; DIV5-7 in DMEM supplemented with 5% horse serum (Thermo Fisher), 10ng/mL IGF-1 (Thermo Fisher), 0.1mM 2-mercaptoethanol, and 2µg/mL doxycycline. See Table S2 for full details of each reagent.

### Lentiviral transduction

TRC lentiviral shRNA plasmids expressing non-targeting (NT) or *SBF1* shRNA (Horizon) were incorporated into lentiviral particles through the University of Michigan Vector Core. Lentiviral lysates were applied to iMN and iSN media on DIV7, or iMuscle media on DIV3.

### Western blot

Cells were pelleted and lysed in RIPA buffer supplemented with protease/phosphatase inhibitor cocktail (Thermo Fisher), followed by sonication and centrifugation at 14000 x g for 10 minutes at 4ºC. Supernatants were transferred to fresh Eppendorf tubes on ice and protein concentrations were determined using BCA assay. Equal protein amounts (20 μg) were resolved on 8-15% SDS-PAGE gels and transferred to PVDF membranes. Membranes were blocked with 3% BSA in TBS-T, incubated overnight with primary antibodies (**Table S1**), washed, and probed with HRP-conjugated secondary antibodies. Bands were detected by enhanced chemiluminescence and imaged using an Amersham Imager 600.

### Immunocytochemistry

Cells were fixed with 4% paraformaldehyde, permeabilized with Triton X-100, and blocked with blocking buffer (3% BSA, 2% fetal calf serum, and 0.1% Triton X-100 in PBS) for 1 hour. Primary antibodies (diluted at 1:100 in blocking buffer) were incubated overnight at 4°C, followed by incubation with Alexa Fluor-conjugated secondary antibodies (diluted at 1:250 in blocking buffer) for 1 hour at room temperature. Nuclei were counterstained with Hoechst 33258 for 5 minutes prior to imaging using an EVOS M5000 fluorescence microscope (Thermo Fisher).

### Propidium Iodide (PI) cell viability assay

Cells were fixed, permeabilized, and incubated in 2X SSC containing DNase-free RNase followed by staining with propidium iodide (Thermo Fisher). After Hoechst counterstaining, images were captured using a fluorescence microscope, PI-positive and Hoescht-positive nuclei in each ROI were quantified using FIJI, and %viability determined by 1 – [(# PI-positive nuclei) / (# Hoescht-positive nuclei)].

### Live-cell fluorescence microscopy

Cells expressing mEGFP-LC3B were imaged on DIV14 (iMNs, iSNs) or DIV7 (iMuscle) using an Oxford Nanoimager (www.oni.bio) at 100X magnification. For drug treatments prior to imaging, cells were treated with 0.25µM VPS34-IN1 or DMSO control for 1 hour prior to imaging. mEGFP-positive puncta were quantified using FIJI (ImageJ).

### Quantification and Statistical Analysis

Independent biological replicates were defined as separate differentiations using distinct media preparations or passages. Image analyses (mEGFP-positive puncta, PI quantification, densitometry of Western blot bands) were performed in FIJI. Data were normalized to loading controls when applicable and expressed as mean ± SEM. Statistical significance of mean differences was assessed using unpaired two-tailed *t*-tests or two-way ANOVA, with *p* < 0.05 considered significant. Data were analyzed and plotted into graphs using Prism (GraphPad).

## RESULTS

We sought to generate a faithful multi-cellular model of the PNS to accurately capture molecular characteristics of CMT4B3 in relevant cell types. To accomplish this, we used an established PiggyBac system^27^ to complete stable genomic knock-in of Tet-ON cassettes (**Fig. 1A**). These cassettes, in the presence of doxycycline, drive expression of transcription factors to direct differentiation of iPSCs into specific cell types of the PNS: *NGN2, ILS1*, and *LHX3* for cholinergic lower motor neurons (iMNs), *BRN3A* and *NGN2* for sensory neurons (iSNs), and *MYOD* and *OCT4* shRNA for skeletal muscle cells (iMuscle)^28-30^ (**Fig. 1A**). We applied this differentiation system in a reference WT iPSC line (KOLF2.1J^31^), and in iPSCs derived from a patient with genetically confirmed CMT4B3 (maternal *SBF1* mutation c.3493_3494dupTA; paternal *SBF1* mutation c.5474_5475delTG) (**Fig. 1B-C**). Within 14 days for iMNs and iSNs, and 7 days for iMuscle, we achieved >99% differentiation of WT and CMT4B3 iPSCs into each PNS cell type (**Fig. 1D-E**). These results confirmed our successful establishment of a multi-cell model of the human peripheral nervous system for studying CMT4B3.

**Figure 1.**
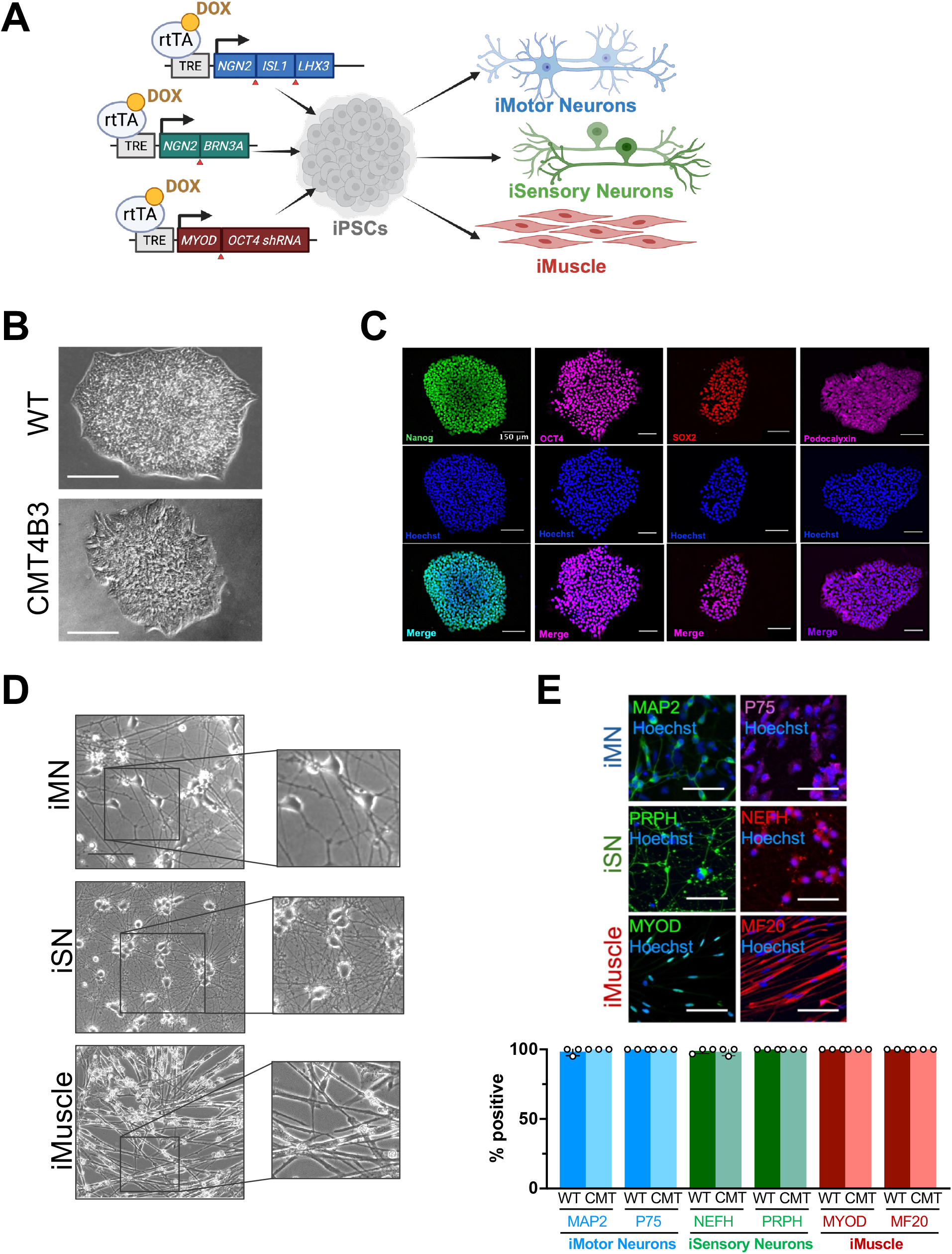
Generation of a multicellular PNS model of CMT4B3. (**A**) Schematic illustrating the PiggyBac-based stable integration of Tet-ON transcription factor cassettes for directed differentiation of iPSCs. (**B**) Representative brightfield microscopy of wild-type (WT) and CMT4B3 patient-derived iPSCs. Scale bar, 0.5mm. (**C**) Immunocytochemical analysis of pluripotency markers in CMT4B3 iPSCs. Scale bar, 150µm (**D**) Representative brightfield microscopy of iPSCs differentiated into induced motor neurons (iMNs), sensory neurons (iSNs), and skeletal muscle cells (iMuscle). Scale bar, 500µm. (**E**) *Top*, immunocytochemical staining of lineage-specific markers in iMNs, iSNs, and iMuscle. Scale bar, 150 μm (iMN, iSN), 300 μm (iMuscle). *Bottom*, quantifications of # marker-positive cells / # Hoescht-positive nuclei.

In almost all cases of CMT4B3, clinical symptoms relate to sensorimotor axon degeneration, without evidence of myopathic features beyond neurogenic atrophy^3,4,6,8-11^. To confirm whether our multi-cell model recapitulated these disease features, we assessed viability in each cell type. Gross morphologic examination revealed marked cell degeneration in CMT4B3 iMNs compared to WT counterparts that became apparent within 3 to 7 days of differentiation (**Fig. 2A**, left column). CMT4B3 iSNs also showed some degree of degeneration compared to WT iSNs, but this was to a less severe extent with greater survival to DIV14 compared to CMT4B3 iMNs (**Fig. 2A**, middle column). CMT4B3 iMuscle cells showed similar survival and morphologic integrity as WT counterparts throughout the entirety of 7 days of differentiation (**Fig. 2A**, right column). To more precisely quantify the extent of these cell type-specific differences in viability among CMT4B3 cells, we next used propidium iodide (PI) staining to label membrane-compromised cells^32^. Strikingly, but consistent with the clinical phenotype of CMT4B3 and our brightfield imaging data, we observed a profound reduction in viability in CMT4B3 iMNs and iSNs (**Fig. 2B-C**). CMT4B3 iMNs showed a 339% increase in PI-positive cells compared to WT iMNs. Similarly, but to a lesser extent, CMT4B3 iSNs showed a 231% increase in PI-positive cells compared to WT iSNs. There was no significant loss of viability in CMT4B3 iMuscle cells compared to WTs. These findings demonstrate a clear, cell type–specific vulnerability to *SBF1*/MTMR5 mutations, similar to what is observed in CMT4B3 patients with peripheral neurons most affected, and motor neurons especially, but muscle cells the least^1-10^.

**Figure 2.**
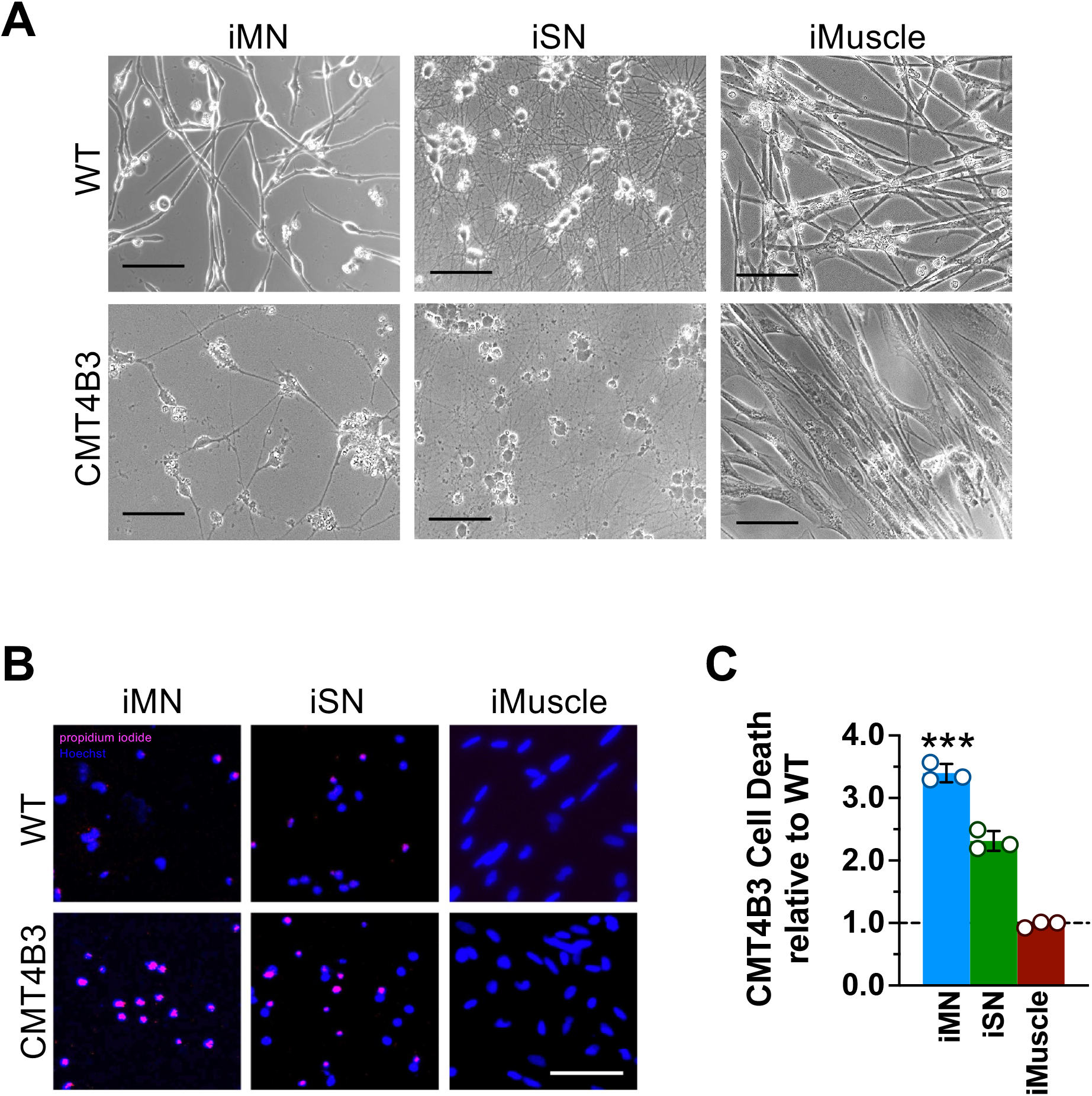
Cell type–specific vulnerability to *SBF1* loss in CMT4B3. (**A**) Representative brightfield images of DIV7 CMT4B3 iMNs, DIV14 iSNs, and DIV7 iMuscle. Scale bar, 500µm. (**B**) Propidium iodide (PI) staining of WT and CMT4B3 iMNs (DIV3), iSNs (DIV14), iMuscle (DIV7). Scale bar, 60µm. (**C**) Relative proportions of cell death in CMT4B4 cells, determined as 1 – (# PI-positive nuclei / # Hoescht-positive nuclei) and normalized to WT cells for each cell type. Data are presented as mean ± SEM (N=3 independent replicates). ***p<0.001, **p<0.01, Student’s t*-*test.

The extent to which myotubularins differ in expression or function in each cell type of PNS is unknown, and this would need to be clarified to fully inform and interpret any disruptions that occur in these proteins due to *SBF1*/MTMR5 mutations. Accordingly, we first assessed the steady state levels of MTMR5 and its functionally related paralogs, MTMR2 and MTMR13, in the WT PNS cells we generated. On Western analysis, we observed that each myotubularin did indeed vary in expression in depending on cell type, with all MTMR5/13 most highly expressed in WT iSNs, but all myotubularins showing lowest expression in isogenic WT iMuscle (**Fig. 3A-D**). In contrast, mutant MTMR5 was significantly attenuated in all CMT4B3 cell types (**Fig. 3A-B**). The most pronounced loss of MTMR5 was seen in iSNs (24% of WT iSN MTMR5) **(Fig. 3B**). MTMR5 loss in CMT4B3 cells correlated with reductions in MTMR2 expression, except for CMT4B3 iMNs, with the greatest loss of MTMR2 in iPSCs (27% of WT iPSC MTMR2) (**Fig. 3A,C**). MTMR13 was reduced but only in CMT4B3 iPSCs and iMuscle (90% of WT iPSC and 65% of WT iMuscle, respectively) (**Fig. 3A, F-G**). These results confirm that *SBF1* mutations are associated with adverse but variable effects on MTMR expression, depending on the cellular context. The greatest reductions occur MTMR2/5 in CMT4B3 cells, but the degree to which the each myotubularin is affected differs by cell type, with CMT4B3 iMuscle the most depleted of all MTMR paralogs.

**Figure 3.**
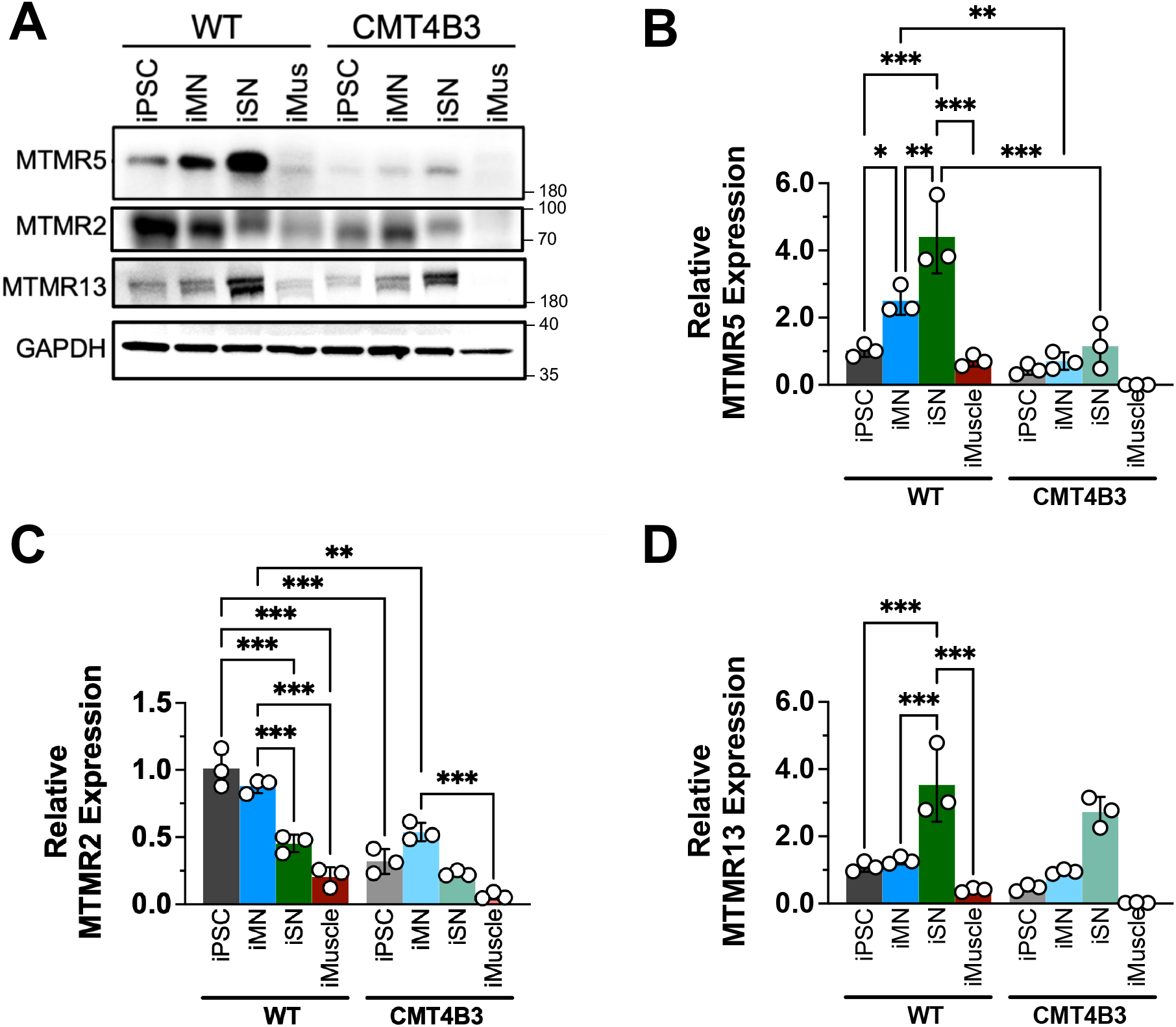
Cell type- and CMT4B3-specific expression of myotubularins PNS cells. (**A**) Representative Western blot of MTMR5, MTMR2, and MTMR13 in WT and CMT4B3 iPSCs, iMNs, iSNs, and iMuscle cells. (**B-D**) Quantifications of relative protein expression normalized to loading control (β-actin) and then normalized to WT iPSC levels. Data are presented as mean ± SEM (N=3 independent replicates). ***p<0.001, **p<0.01, *p<0.05, two-way ANOVA.

Given that MTMR5 and related myotubularins have been implicated in inhibiting autophagy^12,17,21,22^, we wondered whether differences in myotubularin expression impact autophagy in each PNS cell type. To assess this, we analyzed the steady-state levels of two canonical autophagy substrates, LC3-II and p62 (also known as SQSTM1)^33^. At baseline in WT cells, we surprisingly found that iSNs had the relatively highest LC3-II with lowest p62 (**Fig. 4A-C**), suggestive of the highest baseline autophagy among PNS cell types. This contrasted with our expectations because iSNs have the highest expression of MTMR5/13, and thus were expected to have the lowest level of baseline autophagy (**Fig. 4A-C**). We also unexpectedly observed no dramatic changes in LC3-II or p62 turnover in WT iMuscle (**Fig. 4A-C**) despite having lowest expression of MTMR2/5/13 (**Fig. 3A-D**). In comparison to WT cells, CMT4B3 cells showed very slight differences in baseline autophagy that correlated with diminished expression of MTMR2/5/13 (**Fig. 3A-G**), namely mildly greater clearance of p62 in CMT4B3 iPSCs, iMNs, and iMuscle (**Fig. 4A,C-D**). Although several of our observations are discordant with expected alterations in MTMR2/5/13 expression, overall our findings nevertheless highlight that each cellular component of the PNS harbors distinct and non-overlapping patterns of baseline autophagy, and that MTMR loss leads to modest increments in autophagy in cells of the PNS.

**Figure 4.**
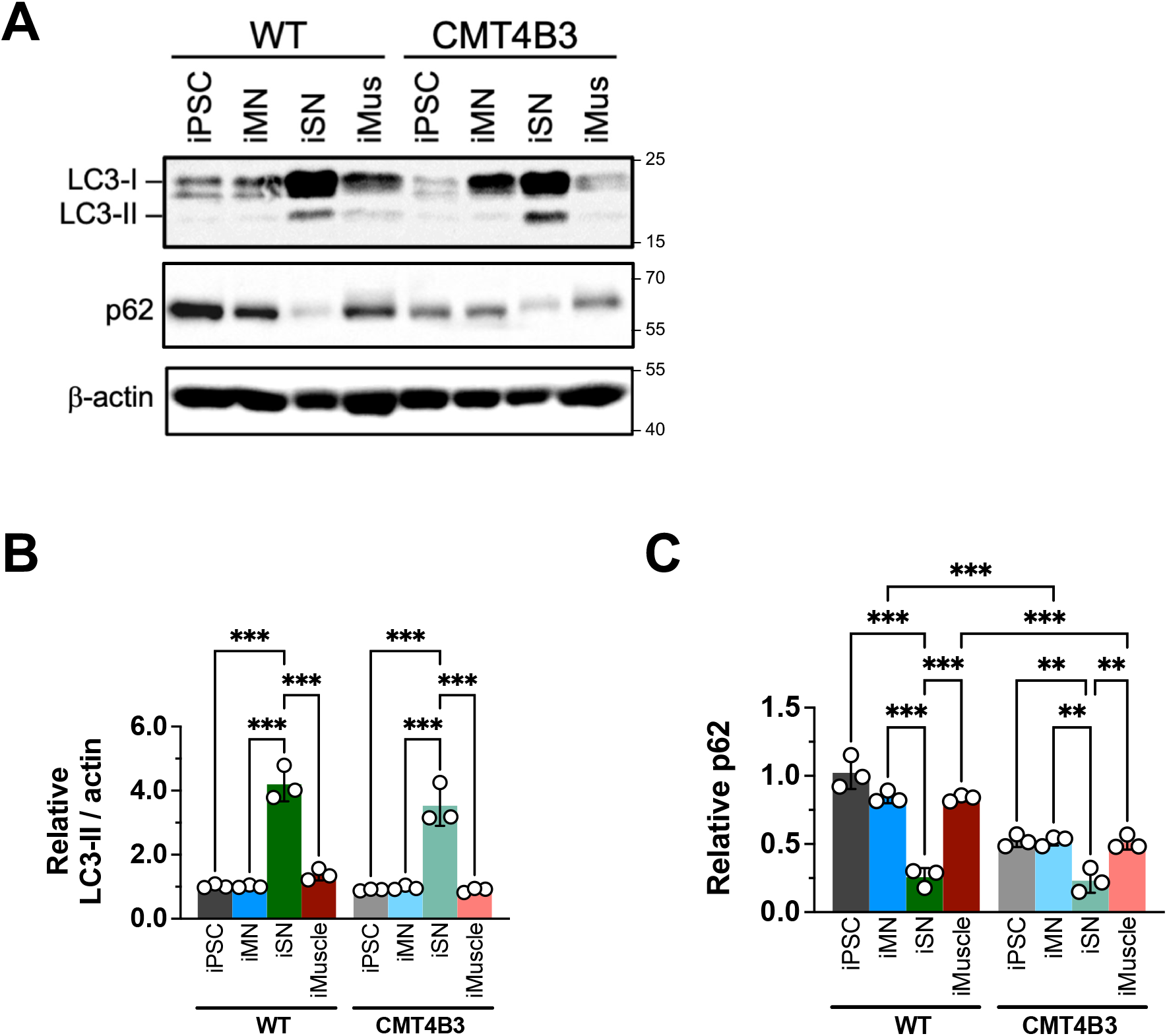
Cell type- and CMT4B3-specific basal autophagy in PNS cells. (**A**) Representative Western blot of LC3-II and p62 in WT and CMT4B3 iPSCs, iMNs, iSNs, and iMuscle cells. (**B-C**) Quantifications of LC3-II and p62 levels normalized to loading control (β-actin), then normalized to WT iPSC levels. Data are presented as mean ± SEM (N=3 independent replicates). ***p<0.001, **p<0.01, *p<0.05, n.s. not significant, two-way ANOVA.

We next asked whether exogenous genetic depletion of MTMR5 was sufficient to mimic molecular phenotypes in a cell type-specific manner, as seen in CMT4B3 patient-derived cells. To do this, we used shRNA-mediated knockdown of *SBF1*/MTMR5 in WT iMNs, iSNs, and iMuscle. These WT PNS cells were differentiated from an iPSC line previously edited by CRISPR/Cas9 to tag LC3B with mEGFP, thereby enabling visualization of autophagosome formation and trafficking in real time by fluorescence microscopy^22^. We then assessed autophagy by live cell imaging and quantified morphologically visible mEGFP-positive vesicles after *SBF1*/MTMR5 knockdown. In neurons, we analyzed axon termini, which are the primary sites of autophagosome biosynthesis^34^, while in muscle cells we focused on perinuclear areas in which omegasomes are enriched^35,36^. Knockdown iMNs had the greatest relative increase in mEGFP-LC3B-positive puncta, with only a mild increase in knockdown iSNs and no difference in knockdown iMuscle (**Fig. 5A-B**). In line with this, knockdown cells showed a distinct pattern of baseline autophagy on Western analysis, contrasting with patient-derived cells: knockdown iMNs had the greatest reductions in LC3-II and p62, with milder reduction of p62 only in iSNs, and milder reduction of LC3-II only in iMuscle (**Fig. 5E-F**). Together, these suggest that acute genetic knockdown of *SBF1*/MTMR5 potentiates autophagy in cell type-dependent manner, with neuronal populations, particularly iMNs, showing the highest disinhibition.

**Figure 5.**
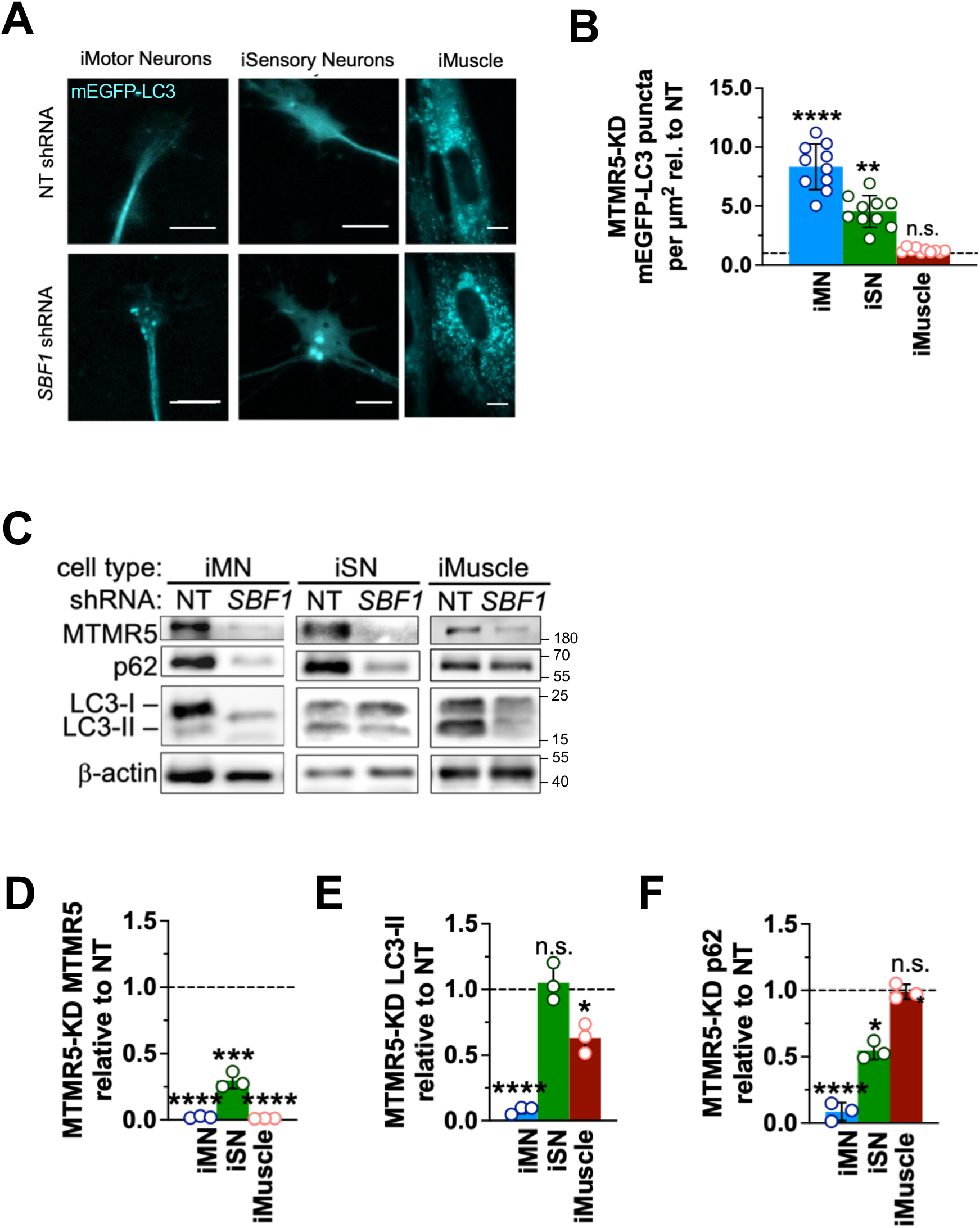
Cell type-specific effects of *SBF1* knockdown on autophagy in PNS cells. (**A**) Western blot of MTMR5 to confirm knockdown efficiency by shRNA, and LC3-II, and p62 to assess basal autophagy, in iMNs, iSNs, and iMuscle cells following shRNA-mediated knockdown of *SBF1*. (**B-D**) Quantifications of MTMR5, LC3-II, and p62 levels in *SBF1* knockdown cells, normalized to loading control (β-actin), then normalized to non-targeted (NT) controls. (E) Representative 100X live-cell epifluorescence microscopy of mEGFP-LC3B puncta in NT and SBF1 knockdown cells. Scale bar, 10µm. (F) Quantifications of mEGFP-positive puncta per µm^2^ in *SBF1* knockdown cells, normalized to NT controls. ****p < 0.001, **p < 0.01, n.s. not significant, Student’s t-test.

Loss of MTMR5 function would be expected to result in unopposed action of the Class III PI3-Kinase (PI3KC3) complex in phosphorylating PtdIns lipids, which may one of the determinative signals driving downstream pathogenic effects in CMT4B3 cells, including autophagy disinhibition^37,38^ (**Fig. 6A**). Based on this, we hypothesized that that pharmacologic inhibition of PI3KC3 might rescue several of the molecular phenotypes we observed in CMT4B3 cells. To test this hypothesis, we used VPS34-IN1, a selective inhibitor of the VPS34 component of PI3KC3 complex^39^ (**Fig. 6A**). In axon termini of iMNs after *SBF1*/MTMR5 knockdown, we observed a significant increase in mEGFP-LC3-positive puncta, but this increase was abrogated after treatment with VPS34-IN1 (**Fig. 6B-C**). Given these results, we next treated CMT4B3 iMNs with VPS34-IN1 and observed significant improvement in cell survival compared to vehicle control (**Fig. 6D-E**). VPS34-IN1 treatment resulted in reduced clearance of LC3-II and p62 (**Fig. 6F-G**) and counteracting the slight autophagy enhancement we observed in CMT4B3 iMNs (**Fig. 4**). However, and surprisingly, we saw no difference in phospho-SGK3, a downstream phospho-substrates of PI3KC3, suggesting that VPS34-IN1 at the concentration used exerted its effects without significant target engagement at PI3KC3. Altogether, our results corroborate that VPS34-IN1 treatment successfully inhibits enhanced autophagy from MTMR5 loss and promotes cell survival in CMT4B3 motor neurons.

**Figure 6.**
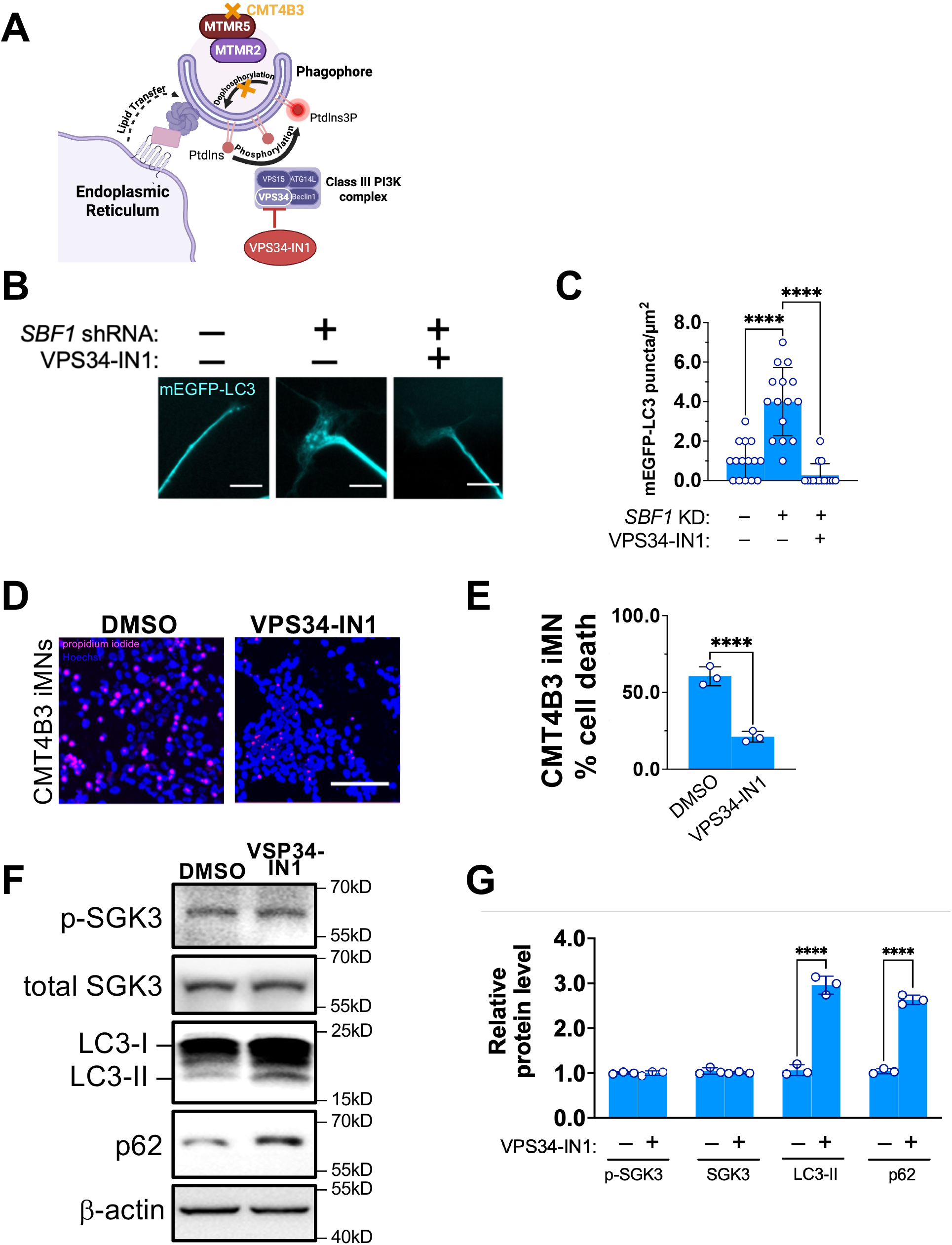
Pharmacologic inhibition of PI3KC3 rescues neurodegeneartion. (**A**) Schematic of phosphatidylinositol (PtdIns) metabolism illustrating the roles of MTMR5 and the Class III PI3-Kinase (VPS34/PI3KC3) complex in phagophore elongation. (**B**) Representative live-cell images of mEGFP-LC3 iMNs following *SBF1*/MTMR5 knockdown and with or without VPS34-IN1 treatment. Scale bar, 10 μm. (**C**) Quantifications of mEGFP-positive puncta per μm^2^ normalized to NT/DMSO-treated control. Data are presented as mean ± SEM (N=3 independent replicates, 5 axons per replicate). ****p<0.0001, two-way ANOVA. (**D**) Propidium iodide (PI) staining of CMT4B3 iMNs treated with and without VPS34-IN1. Scale bar, 60µm. (**E**) Quantifications of cell death in CMT4B3 iMNs treated with and without VPS34-IN1, determined as 1 – (# PI-positive nuclei / # Hoescht-positive nuclei). Data are presented as mean ± SEM (N=3 independent replicates, 100 nuclei scored per replicate). ****p<0.0001, Student’s t-test. (**F**) Representative Western blot of PI3KC3 pathway phospho-substrate SGK3, LC3-II, and p62 in iMNs with and without VPS34-IN1 treatment. (**G**) Quantifications of band intensities in (F) normalized to loading control (β-actin), then normalized to DMSO-treated cells. Data are presented as mean ± SEM (N=3 independent replicates). ***p < 0.001, **p < 0.01, *p < 0.05, Student’s t-test.

## DISCUSSION

In this study, we established the first human peripheral nerve cell model of MTMR5 biology and CMT4B3. Our model system successfully generated essentially pure populations of human motor neurons, sensory neurons, and skeletal muscle (**Fig. 1D-F**). This aspect is especially important for overcoming barriers imposed on other model systems of CMT4B3. With the composition of our model system consisting entirely human peripheral nerve cells, it most closely models the target cell types predominantly affected by the disease. In contrast, prior work has investigated MTMR5 function in COS-1 cells^14^, HEK293 cells^14^, HeLa cells^15^, human fibroblasts^20,23^, undifferentiated iPSCs^40^, and non-PNS neurons and glial^22^ cells, as well as zebrafish^41^ and mice^25,42^. Crucial mechanistic insights about MTMR5 have been gleaned from these studies, such as interactions with MTMR2^14^, subcellular distribution^15^, facilitating PtdIns metabolism and suppressing autophagy^14,22^, and roles in radial axon sorting and spermatogenesis^2543^. However, since these studies do not directly assess MTMR5 in human peripheral nerve cells, the expression levels and activity of MTMR5 reported might be insufficiently reflective of human MTMR5 biology and thus limits translating such insights into understanding CMT4B3 pathogenesis in human patients. Several observations are in line with this: murine and zebrafish *Sbf1* loss has minimal neuromuscular pathology that is not commensurate with the severity of human phenotypes^41^, and male human patients do not harbor the infertility phenotype seen in KO mice^25,42^. Although our present study has revealed several discoveries about MTMR5 biology, such as cell type-specific expression patterns (**Fig. 3**), there are still several areas to further define MTMR5 function in the PNS and relevant CMT4B3 pathobiology. For example, corroborative studies to establish *SBF1* mutation-dependent effects should use CMT4B3 patient cells that have been CRISPR-corrected to WT. In addition, our PNS system lacks interrogation of MTMR5 biology in other aspects of the PNS, specifically in Schwann cells and in the 3-D complexity of the multicellular neuromuscular apparatus of the peripheral nerve (i.e., nerves with myelination, neuromuscular junctions, and muscle contiguity). These gaps should be addressed in future studies to better understand cell-specific and non-cell autonomous aspects of disease.

Our study uncovered several important observations about myotubularin expression. The steady-state levels of MTMR5 in WT cells confirm that MTMR5 is neuronally enriched (i.e., in iMNs and iSNs; **Fig. 3A-B**), as previously described^11,12^. These observations are an important starting point and human MTMR expression should be corroborated in samples of each cell and tissue type from human patients. Second, MTMR5 expression level is attenuated in all cell types when expressing CMT4B3-related mutations (**Fig. 3A-B**). Although the exact mechanism connecting the mutations with this loss of expression is not elucidated in the present study, we highly suspect this is related to nonsense-mediated decay (NMD)^43^ given the introduction of premature termination codons by both mutations (**Fig. 1**). If so, candidate therapy strategies include inhibiting NMD^44,45^ or targeting alternative splicing^46^ to restore MTMR5 expression, depending on the extent to which increased mutant MTMR5 protein may cause toxic dominant negative effects, but these pathobiological details require further investigation. Separately, in CMT4B3 cells and with loss of MTMR5 expression, there was concomitant loss of MTMR2, but not MTMR13, in multiple cell types (**Fig. 3A,C**). This finding is in line with the regulation of MTMR2 stability by MTMR5^14,15^ and also suggests against MTMR5 having a similar role for MTMR13, overall corroborating that inter-myotubularin regulatory activities are unique to each myotubularin^15^. Lastly, given that MTMR2 is the principal effector of the MTMR5-MTMR2 axis^14^, another candidate strategy for treating CMT4B3 is to compensate for MTMR2 loss using gene therapies that restore or overexpress MTMR2, but this also requires further investigation.

One surprising result of our study is that autophagy was not robustly altered by loss of MTMR5 and MTMR2 in neurons (**Fig. 4**). Given the putative inhibitory effects of both myotubularins on autophagic flux^14,22^, and that *SBF1* knockdown in WT iMNs and iSNs seemed to dramatically disinhibit autophagy (**Fig. 5**), we expected that reduced expression levels of these myotubularins would be inversely correlated with autophagy activity. Nonetheless, our findings raise several intriguing possibilities about MTMR5 function. MTMR2 or MTMR5 loss in CMT4B3 may have only minor or no determinative roles in dysregulating autophagy to cause toxicity in peripheral neurons compared to other cell types (**Fig. 2**). Acute loss of MTMR5 by shRNA knockdown may have discordant effects compared to chronic loss due to CMT4B3 mutations, wherein initially or in the short-term a reduction of MTMR5 effectively enhances autophagy (consistent with our prior work^22^), but longer term absence of MTMR5 may lead to compensatory mechanisms that re-balance autophagy, such as by regulating expression of other myotubularins^41^. Other myotubularins, especially the 11 that have not been assessed in this study, may also have more potent regulatory effects on autophagy in peripheral neurons^15^. Separately, MTMR2/5 loss in CMT4B3 may exert toxic effects through loss of other functions such as catalyzing GDP/GTP exchange^47^ or cellular growth^48^. To underscore evidence pointing to functional nuances of each MTMR in different cell contexts, CMT4B3 sensory neurons had less severe cell death despite greater expression loss of MTMR5 compared to isogenic motor neurons (**Fig. 2**). These data indicate that CMT4B3 pathogenesis is not determined solely by the magnitude of MTMR5 protein loss, but vulnerability may emerge from the intersection of depletion severity and cell type–specific requirements for MTMR function (with motor neurons emerging as the most sensitive to disruption). Overall, our findings highlight that each cellular component of the PNS harbors distinct and non-overlapping patterns of myotubularin expression, autophagy function, and cell viability. Given the discordance between MTMR expression and autophagic flux seen in CMT4B3 cells, future studies will need to clarify the salient pathomechanisms of MTMR5 loss and assess how the global molecular landscape downstream of MTMR5 is altered in each peripheral nerve cell type.

Our multi-cell model of CMT4B3 showed cell death primarily in iMNs and iSNs (**Fig. 2A-B**), and this observation is critically important for several reasons. First, these results confirm the biological fidelity of our model because cell degeneration restricted to sensorimotor neurons correlates with the peripheral axonopathic phenotype seen in CMT4B3 patients (including the patient from which our mutant iPSCs were derived), and our model did not produce an unexpected myopathic phenotype. Second, and more crucially, this is a well-defined and disease-relevant phenotype that is targetable for rescue in therapeutic testing. Successful rescue of this phenotype in our model would greatly enhance the clinical trial readiness of therapy candidates because reversal of peripheral neuron death is expected to be of immense clinically meaningful benefit, given the severely disabling manifestations of progressive limb weakness and wheelchair dependence in CMT4B3 patients.

Notably, our cellular platform has a high degree of scalability with nearly limitless amounts of PNS cells able to be generated from our self-renewing iPSC lines. Coupling this scalability with the essential disease feature of peripheral nerve degeneration, our cellular platform can thus faithfully serve to validate and accelerate candidate therapies for preclinical testing in a disease-specific and high-throughput manner. These features of our model system should be leveraged in tandem with the most exciting result from our study, which implicates VPS34 inhibition as a viable therapeutic intervention to significantly improved peripheral neuron survival (**Fig. 6A-D**). Mechanistically, confirmation of curtailing autophagy disinhibition implicates at least some role of autophagy dysregulation^39^ in mediating neurotoxicity (**Fig. 6E-F**), but this may be independent of PtdIns regulation given that altering PI3KC activity was not necessary to mediate the rescue effect of VPS34-IN1 (**Fig. 4**). Of not, a key barrier to translational application of VPS34-IN1 is lack of FDA approval for clinical use in human patients. However, other analogs that have similar effects are FDA-approved, including compounds in a library of autophagy regulators for motor neurons^49^ that we previously validated, and which now warrant similar testing to facilitate clinical trial translation for CMT4B3. Altogether, our innovative and powerful cellular model of CMT4B3 has established critical and foundational groundwork for defining novel mechanisms of pathogenesis and refining molecular strategies to accelerate therapeutic development and interventions for this devastating and incurable disease.

## ACKNOWLEDGEMENTS

We thank the Schultz family for their generosity, collaboration, coordination, and support in securing CMT4B3 patient-derived iPSCs and guiding research design and experimental prioritization. Figures 1A and 7A were created using BioRender (www.biorender.com). This research was supported by a research award from The CMT4B3 Foundation (Grant No. 145128, RCC 1708211501).

