## Supplemental Files for "Defining and rescuing pathomechanisms of myotubularin and autophagy disruption in a novel human cell model of Charcot-Marie-Tooth Type 4B3"

### Supplementary Tables

#### Table 1

| Antibody | Dilutions | Vendor | Catalog No. |
|---|---|---|---|
| Primary Antibodies | | | |
| Rabbit polyclonal anti-MTMR5 | 1:1000 | Abcam | ab181106 |
| Rabbit polyclonal anti-LC3B | 1:1000 | Cell Signaling | 2775 |
| Mouse monoclonal anti-p62 | 1:2000 | Novus Biological | H00008878-M01 |
| Rabbit polyclonal anti-β-Actin | 1:5000 | Cell Signaling | 4967S |
| Mouse polyclonal anti-GAPDH | 1:2000 | ProteinTech | 60004-1-Ig |
| Rabbit polyclonal anti-MAP2 | 1:400 | ProteinTech | 17490-1-AP |
| Rabbit polyclonal anti-beta III tubulin | 1:400 | Abcam | ab18207 |
| Rabbit polyclonal anti-CHAT | 1:400 | ProteinTech | 20747-1-AP |
| Rabbit polyclonal anti-Peripherin | 1:400 | ProteinTech | 17399-1-AP |
| Rabbit monoclonal anti-NeuN | 1:400 | Abcam | ab177487 |
| Rabbit polyclonal anti-NEFH (NF-H/NF-200) | 1:400 | ProteinTech | 18934-1-AP |
| Rabbit polyclonal anti -MyoD1 | 1:400 | ProteinTech | 18943-1-AP |
| Mouse polyclonal anti -MF20 | 1:400 | R&D systems | MAB4470SP |
| Rabbit polyclonal anti -P75 NTR | 1:400 | ProteinTech | 55014-1-AP |
| Rabbit polyclonal anti-Oct4 | 1:400 | ProteinTech | 11263-1-AP |
| Rabbit polyclonal anti-NANOG | 1:400 | ProteinTech | 14295-1-AP |
| Rabbit polyclonal anti-Sox2 | 1:400 | ProteinTech | 11064-1-AP |
| Rabbit polyclonal anti-Podocalyxin | 1:400 | ProteinTech | 18150-1-AP |
| **Secondary Antibodies for Western Blots** | | | |
| Rabbit IgG (H+L) Highly Cross-Adsorbed Secondary Antibody | 1:10000 | ThermoFisher | A16110 |
| Mouse IgG (H+L) Highly Cross-Adsorbed Secondary Antibody | 1:10000 | ThermoFisher | A16078 |
| **Secondary Antibodies and Dyes for ICC/ PI** | | | |
| IgG (H+L) highly cross-adsorbed AlexaFluor 488 Goat anti-Rabbit | 1:250 | Invitrogen | A32731 |
| IgG (H+L) highly cross-adsorbed AlexaFluor 488 Goat anti-Mouse | 1:250 | Invitrogen | A32723TR |
| IgG (H+L) highly cross-adsorbed AlexaFluor 647 Goat anti-rabbit | 1:250 | ThermoFisher | A32733TR |
| IgG (H+L) highly cross-adsorbed AlexaFluor 568 Goat anti-rabbit | 1:250 | ThermoFisher | A11011 |
| Hoechst 33258 | 1:5000 | Sigma | 94403-1ML |

#### Table 2

| Media supplements | Vendor | Catalog No. |
|---|---|---|
| DMEM/F12, HEPES | ThermoFisher Scientific | 11330057 |
| MEM alpha, no phenol red | Invitrogen | 41061029 |
| Opti-MEM | Gibco | 31985-070 |
| Neurobasal -A Medium, no phenol red | ThermoFisher Scientific | 12349015 |
| BrainPhys Without Phenol Red | StemCell Technology | 05791 |
| FluoroBrite DMEM | ThermoFisher Scientific | A1896701 |
| L-Ascorbic acid | Millipore-Sigma | A8960 |

| | | |
|---|---|---|
| Sodium Selenite | Millipore-Sigma | S5261 |
| Transferrin | Millipore-Sigma | T3705 |
| Insulin | Biogems | 10-365-1G |
| FGF2 | PeproTech | 100-18B |
| TGF- b | PeproTech | 100-21 |
| N2 supplement | ThermoFisher Scientific | 17502048 |
| MEM Non-essential Amino acids (NEAA) solution (100x) | Millipore-Sigma | M7145-100ML |
| Recombinant human/murine/rat BDNF | PeproTech | 450-02-10ug |
| Compound E | BioGems | 2091746 |
| Human NT-3 Recombinant Protein | PeproTech | 450-03-10ug |
| Laminin | Millipore-Sigma | L2020-1MG |
| Glutamax | ThermoFisher Scientific | 35050061 |
| CultureOne (100x) | ThermoFisher Scientific | A3320201 |
| 5% Knockout serum replacement | ThermoFisher Scientific | 10828028 |
| Sodium pyruvate solution | Sigma Aldrich | S8636 |
| 100x Penicillin- Streptomycin | Millipore Sigma | P0781 |
| 2- Mercaptoethanol | VWR LifeScience | 97064-588 |
| 5% Horse serum | ThermoFisher Scientific | 16050130 |
| Recombinant Human IGF-I/IGF-1 Protein | R&D Systems | 291-G1 |
| Y-27632 Dihydrochloride (ROCKi) | Fisher (1012911) | 501012280 |
| Doxycycline | Millipore-Sigma | 1225984 |
